# 2,6-diaminopurine enhances Aqp4 stop codon readthrough and *AQP4* perivascular localization in the brain

**DOI:** 10.64898/2026.07.27.741080

**Authors:** Sadan Dahal, Ezzeldin N Elhawary, Prarthana Suresh, Zhan Shi, Hamza Lalami, Alyssa M Le, Darshan Sapkota

## Abstract

In the brain, the water channel Aquaporin 4 (AQP4) is largely restricted to astrocytes, enriched at perivascular astrocytic processes, and involved in fluid balance and neurological disease. The perivascular pool is functionally unique as it is depolarized in diseases, targeted by neuromyelitis optica autoantibodies, and implicated in the clearance of brain metabolites such as amyloid beta. The perivascular AQP4 was recently shown to be AQP4X, an extended isoform arising from readthrough, where approximately 20% of translating ribosomes continue through the *Aqp4* stop codon. Because *Aqp4* terminates with a UGA, here we tested whether *Aqp4* readthrough can be enhanced by 2,6-diaminopurine, a purine analog known to promote UGA decoding. Using dual luciferase and immunoblotting for AQP4X in cultured cells, we show that the drug promotes *Aqp4* readthrough and increases AQP4X levels. For *in vivo* validation, we use the wild-type mouse as well as a genetically engineered mouse with a stop-to-sense mutation allowing *Aqp4* readthrough at 100%. 2,6-diaminopurine elevates perivascular AQP4X levels by approximately 10% after 20 hours of single intracranial injection. It fails to enhance AQP4X levels in animals with stop-to-sense mutation, suggesting its action involves *Aqp4* readthrough. *Aqp4* readthrough can be a pharmacological target for modulating perivascular AQP4X in neurological disease models.

## INTRODUCTION

The water channel protein Aquaporin 4 (AQP4) plays critical neurological roles due to its abundance and distribution in the central nervous system. Its function in fluid and ion homeostasis is manifest in *Aqp4-/-* mice. At baseline or upon experimental challenges, *Aqp4-/-* mice exhibit altered synaptic plasticity, neuronal activity, seizure dynamics, extracellular space volume, and edema vulnerability^1–4^. AQP4 is found predominantly at the cell membrane of astrocytes, with strong enrichment at specialized processes called astrocyte endfeet that wrap the blood-brain barrier^5^. The perivascular pool of AQP4 is noteworthy for several reasons. For one, it is depolarized in aging, neurodegeneration, epilepsy and glioma, and its repolarization is thought to have a translational value^6–10^. Additionally, the perivascular AQP4 is implicated in removing brain metabolites through the exchange of cerebrospinal fluid and interstitial fluid at perivascular spaces. In *Aqp4* global knockout mice, the clearance of toxic proteins such as amyloid beta and tau from the brain parenchyma is compromised, suggesting a role for overall AQP4^11,12^. The same deficit surfaces in mice that lack perivascular AQP4 or the scaffold protein alpha-syntrophin responsible for docking AQP4 at the perivascular region, ascribing the clearance mechanism to the perivascular pool of AQP4^13,14^. Finally, the perivascular AQP4 is preferentially targeted by pathogenic antibodies in the autoimmune demyelinating disease neuromyelitis optica^15^. Thus, targeting perivascular AQP4 may be one way to regulate AQP4 function and modify AQP4-associated diseases.

Unfortunately, no genetic handle existed for selectively modulating perivascular AQP4 until recently, when we and others demonstrated that it was a distinct molecular isoform generated by stop codon readthrough^16,17^. In readthrough, some of the translating ribosomes continue through the stop codon until a second stop in the 3’ untranslated region. Approximately 20% of ribosomes exhibit this on *Aqp4* and extend the polypeptide chain by 28 extra amino acids at the C-terminus. Two classical isoforms of AQP4, namely M1 and M23, which arise from two different initiation codons, have been long known^18^. Readthrough results in M1X and M23X, collectively termed AQP4X. In Aqp4NoX mice, which have two extra stop codons immediately behind the annotated stop codon, perivascular AQP4 is selectively lost while the parenchymal pool is intact^14^. Conversely, in Aqp4AllX mice with the annotated stop mutated to a sense to allow constitutive readthrough, perivascular AQP4 is enhanced^19^. Readthrough, therefore, affords a means to modulate perivascular AQP4.

This genetic insight has enabled new functional inquiries into perivascular AQP4. We found that Aqp4NoX mice have impaired clearance of amyloid-beta^14^, possibly due to reduced perivascular clearance. Others have reported that these mice also exhibit diminished reactivity to neuromyelitis optica autoantibody, as the target perivascular pool is depleted^20^. It is possible that Aqp4AllX mice may also show a blunted autoantibody binding because although they are rich in perivascular AQP4, they have reduced levels of AQP4 supramolecular complexes, which are actual epitope complexes^19^. These observations from genetic models position AQP4X as a molecular lever for intervening in diseases. However, how can we leverage *Aqp4* readthrough for this purpose? Using a reporter assay to screen small molecules, we previously identified candidates that enhance the phenomenon *in vitro*^14^. The compounds, nevertheless, were not tested for their ability to enhance AQP4X *in vivo*, nor were they clinically approved ones.

The current study sought to evaluate the *in vivo* efficacy of 2,6-diaminopurine (DAP), a natural adenine analog known to correct UGA nonsense mutations that lead to truncated proteins and non-sense mediated decay of mRNAs^21–23^. The drug is known to promote the translation of impacted mRNAs into full-length proteins. Notably, the annotated stop codon of *Aqp4* is a UGA. Using reporter and immunoblot assays in cell cultures, we demonstrate that DAP robustly promotes *Aqp4* readthrough and AQP4X expression. Further, DAP treatment in wild-type mice leads to an increase in the perivascular localization of AQP4X. In contrast, the drug is no longer effective in Aqp4AllX mice which lack the annotated stop codon, indicating it functions by enhancing *Aqp4* readthrough. Thus, we identify perivascular localization of AQP4 as a druggable process and provide a proof of concept that a readthrough enhancing drug can be leveraged to selectively promote this process.

## RESULTS

### DAP enhances *Aqp4* readthrough in cell-based assays

To assess the effect of DAP on *Aqp4* readthrough, we employed a dual-luciferase reporter assay as described previously^24^. The reporter vector consists of Renilla luciferase (RL) and Firefly luciferase (FL), between which an *Aqp4* test cassette is cloned in frame, such that the former is upstream and expressed constitutively, while the latter is expressed only if ribosomes read past the UGA stop codon of *Aqp4*. In a control cassette, UGA is mutated to UGG so that FL is also constitutive. Readthrough rate is obtained by normalizing the FL/RL ratio of the test by that of the control **(Fig. 1A)**. Upon transfecting N2a cells, we observed a rate of 15% for *Aqp4*, and a 24-hour treatment with G418, an antibiotic known to enhance readthrough, increased this rate five-fold **(Fig. 1B)**. These values, both baseline and G418-enhanced, align with prior reports^16,17,25^ and lend credibility to the dual-luciferase approach. Treatment with DAP for 24 hours resulted in a clear dose-dependent increase in *Aqp4* readthrough **(Fig. 1B)**, with an EC_50_ (half-maximal effective concentration) of 27 µM **(Fig. 1C)**. Thus, DAP is a robust enhancer of *Aqp4* readthrough in cultured cells.

**Figure 1.**
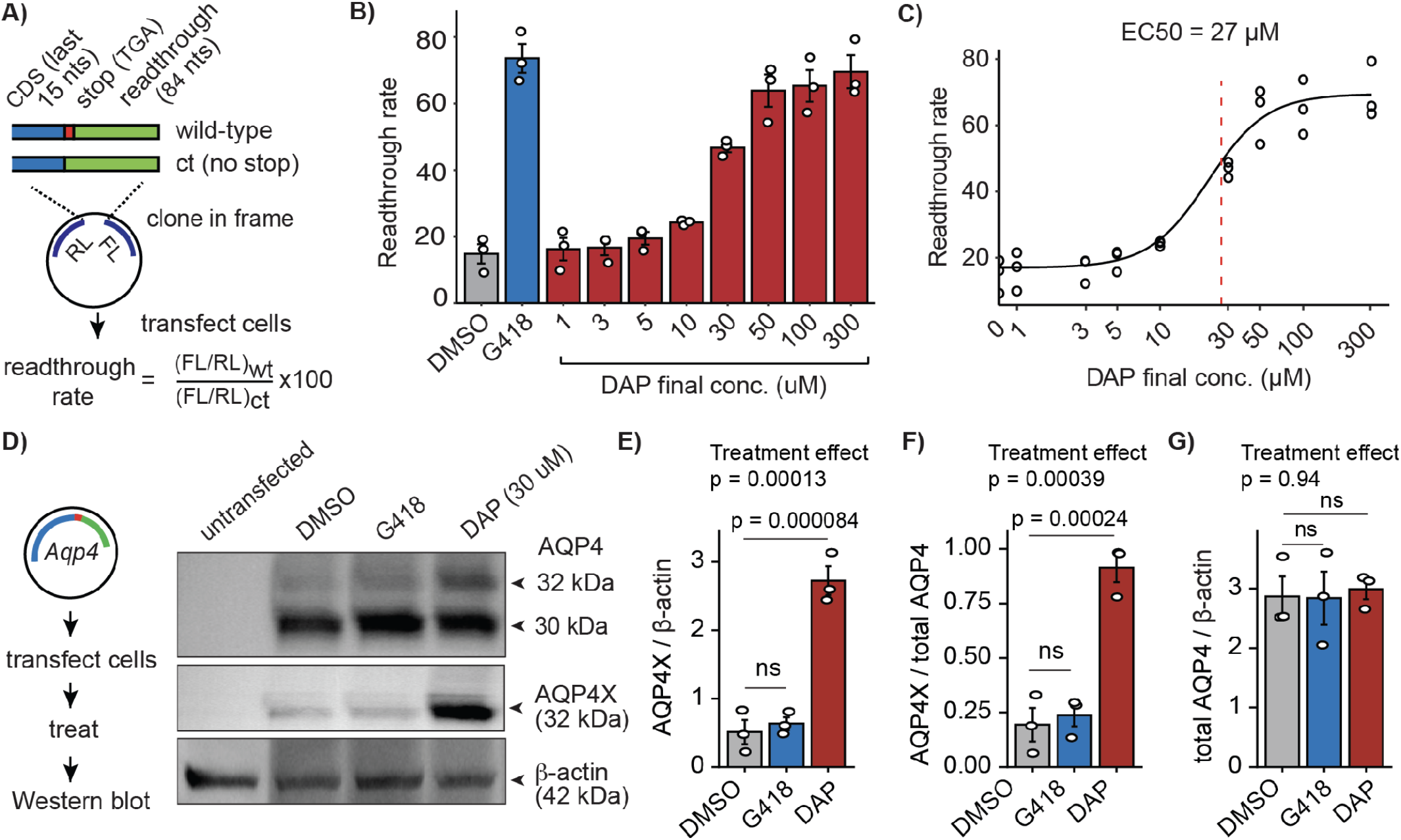
DAP enhances *Aqp4* readthrough in reporter and cell culture assays. **(A)** Schematic of dual-luciferase assay. Indicated cassettes were cloned in-frame such that Renilla was constitutive, but Firefly was reliant on stop codon readthrough. *Aqp4* readthrough rate was quantified after transfecting N2a cells. **(B)** Bar plot shows DAP dose-dependent increase in readthrough in transfected cells after 24 h of treatment. G418 is positive control drug. N = 3 biological replicates. Circles indicate mean of three technical replicates, and error bars indicate standard errors of mean. **(C)** DAP effect is replotted to calculate half maximal effective concentration (EC50). **(D)** Western blot following N2a cell transfection with an *Aqp4* expression plasmid and 24 h treatment with drugs. **(E-G)** Quantification of D. One-way ANOVA, F[2,6] = 55.34(E), 37.75 (F), 0.056 (G). P values at the top show the overall effect of treatment. P values for the effects of G418 and DAP were obtained with Dunnett’s multiple comparison. N = 3 biological replicates, indicated by circles. Error bars indicate standard errors of mean. CDS: coding sequence; nts: nucleotides; RL: Renilla luciferase; FL: Firefly luciferase.

Some small molecules modulate luciferase enzyme activity or stability and hence alter the FL/RL ratio, spuriously appearing as readthrough regulators^26,27^. A genuine candidate should increase the level of a readthrough product. To validate DAP as a bona fide regulator, we sought to quantify AQP4X directly. A vector containing ‘full-length *Aqp4* cDNA-stop-readthrough region’ was transfected into N2a cells, which are neuroblastoma cells that do not express AQP4 endogenously. Cells were treated with G418 (500 μg/mL) or DAP (30 µM, ~ EC50 dose) for 24 hours, and total AQP4 and AQP4X were quantified using Western blot. Anti-AQP4 antibody resolved two bands corresponding to normal-length AQP4 (30 kDa) and higher molecular weight AQP4X (32 kDa), whereas anti-AQP4X antibody solely recognized the latter **(Fig. 1D)**. When compared to DMSO, G418 did not enhance AQP4X levels appreciably in this assay **(Figs. 1E,F)**. DAP, however, upregulated AQP4X five-fold, and this was true relative to the actin loading control as well as total AQP4 level **(Figs. 1E,F)**. DAP treatment also enhanced the intensity of the upper band corresponding to AQP4X in the anti-AQP4 blot **(Fig. 1D)**, although this isoform was quantified only using the anti-AQP4X blot. Consistent with a selective effect on readthrough, the total AQP4 levels remained unchanged in response to treatments **(Fig. 1G)**. These orthogonal immunoblot results are consistent with the dual luciferase findings, establishing DAP as a genuine enhancer of *Aqp4* readthrough *in vitro*.

### DAP enhances perivascular localization of AQP4 *in vivo*

We then asked whether DAP could elevate AQP4X abundance *in vivo*. DAP is quickly eliminated through urine and retained in different tissues and becomes undetectable in the plasma and the CNS within 2 hours after intravenous or oral administration^22^. In line with this uncertainty around CNS bioavailability after a systemic exposure, in our pilot experiments, intravenous injection of DAP (8.1 mg/kg, twice daily for 5 days) did not alter brain AQP4X levels. We therefore resorted to direct intracranial administration for potential proof-of-concept findings. We started with wild-type mice, in which the brain exhibits *Aqp4* readthrough at about 20% at baseline^16^. DAP (300 µM, total volume 2 μL) was stereotaxically injected into the dorsal hippocampus of one hemisphere. This dose was selected based on *in vitro* findings showing robust readthrough enhancement without overt cytotoxicity. An equal volume of PBS was injected into the contralateral hemisphere. Twenty hours later, AQP4X was quantified using immunofluorescence staining. Analysis focused on the hippocampus surrounding the injection site and adjacent cortical regions **(Fig. 2A)**. Compared to PBS-injected hemispheres, DAP-injected hemispheres displayed >10% increase in AQP4X immunoreactivity in both cortex and hippocampus **(Fig. 2B)**. We note that this increase, while modest in magnitude, was elicited by a single DAP injection.

**Figure 2.**
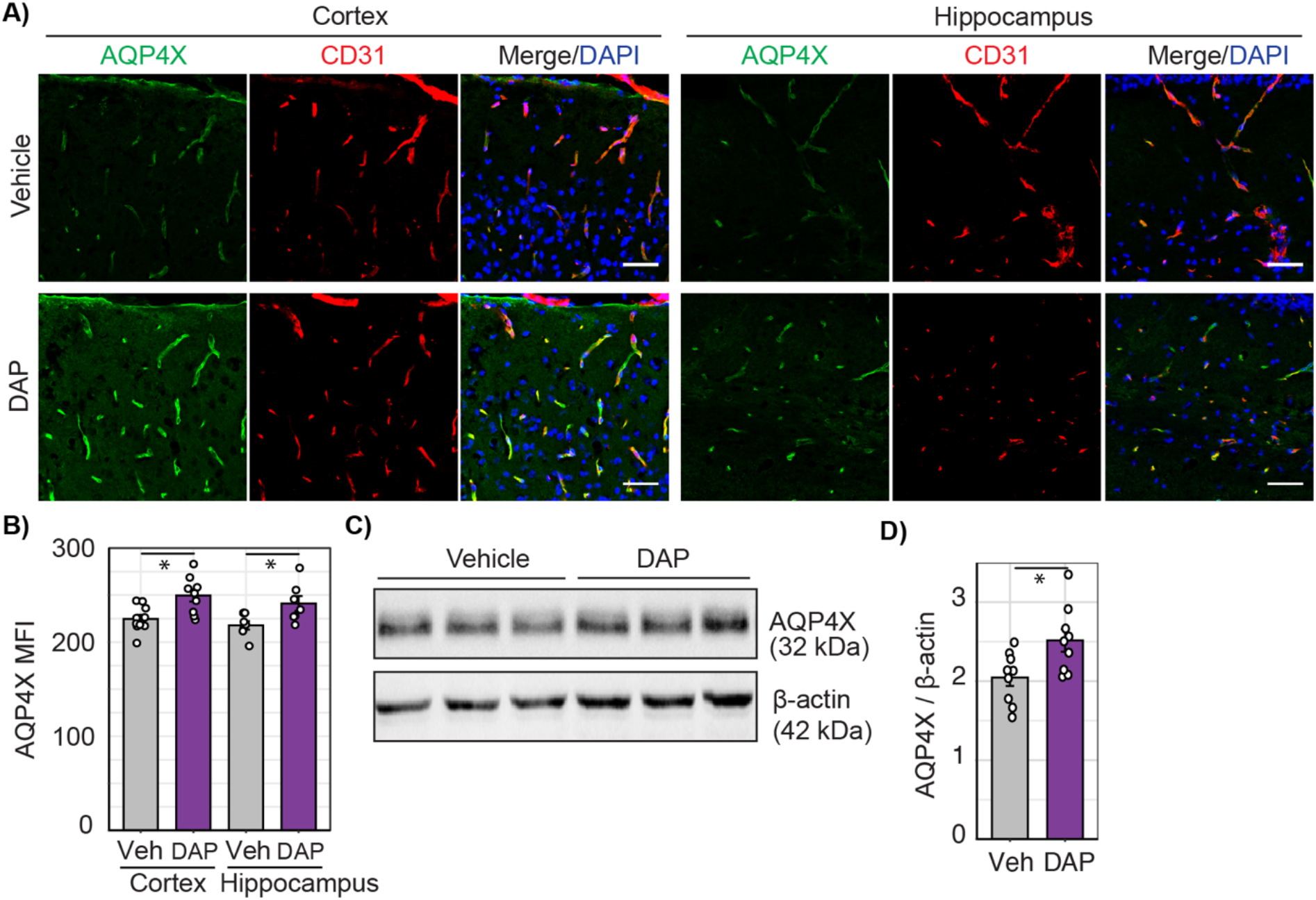
DAP enhances perivascular localization of AQP4 *in vivo*. **(A)** Wild-type mouse brain hemispheres injected with DAP (300 uM) or vehicle, and AQP4X and CD31 visualized in the cortex and hippocampus at or near the injection site using immunofluorescence staining after 20 hours. DAPI marks nuclei. Scale bars = 50 uM **(B)** Quantification of A for mean fluorescence intensity (MFI) of AQP4X. Student’s *t*-test, N = 3 hemispheres per treatment group. Error bars = standard errors of mean. Circles represent technical replicates. ∗p ≤ 0.05 **(C)** Wild-type mouse brain hemispheres injected with DAP (450 uM) or vehicle and injected regions assayed with Western blot after 20 hours. **(D)** Quantification of C. Student’s *t*-test, N = 9 hemispheres per treatment group. Error bars = standard errors of mean. Circles represent biological replicates. ∗p ≤ 0.05.

To independently confirm the in vivo effect of DAP, we attempted immunoblotting for AQP4X in the two hemispheres. In pilot experiments, single intracranial injections indicated a trend but failed to produce statistically significant results. The local effect of DAP apparent with immunofluorescence staining could have been obscured in immunoblotting which requires a substantially larger tissue mass. We therefore injected two adjacent sites per hemisphere, while also increasing the volume of DAP and PBS from 2 µL to 3 µL. Subsequent quantification revealed 14% increase in AQP4X levels in DAP-injected hemispheres compared to PBS **(Figs. 2C, D)**. Taken together, immunofluorescence and immunoblot results show that DAP enhances AQP4X abundance.

### DAP elevates AQP4X levels via translational readthrough

Although cell culture and *in vivo* results showcase DAP’s ability to upregulate AQP4X, they do not quite confirm stop codon readthrough as the underlying mechanism. Since normal-length AQP4 and extended AQP4X are translated from the same mRNA, regulation of transcription or mRNA stability is irrelevant. Given that AQP4X is upregulated but AQP4 is unaltered (Figs. 1F, G), overall increase in translation is also unlikely to account for DAP’s effect. Increased stabilization of AQP4X, however, remains a possible mechanism. To test this, we utilized the Aqp4AllX mouse in which the *Aqp4* stop codon is mutated to a sense codon to allow constitutive expression of AQP4X. We reasoned that DAP should cease to upregulate AQP4X levels in Aqp4AllX mice if it enhances readthrough, but not if it acts by stabilizing AQP4X. DAP (300 µM, 2 µL) or PBS was stereotaxically injected in the dorsal hippocampus of Aqp4AllX homozygous mice at single sites, and AQP4X was quantified by immunofluorescence after 20 hours as above. Note that the half-life of AQP4 is about 24 hours^28^, and therefore increased stability of AQP4X should be appreciable in 20 hours. Indeed, DAP failed to raise AQP4X levels compared to PBS **(Fig. 3A, B)**. By arguing against altered stability as the primary mechanism, these data from Aqp4AllX mice, together with cell culture results, strongly suggest that DAP augments AQP4X production through translational readthrough.

**Figure 3.**
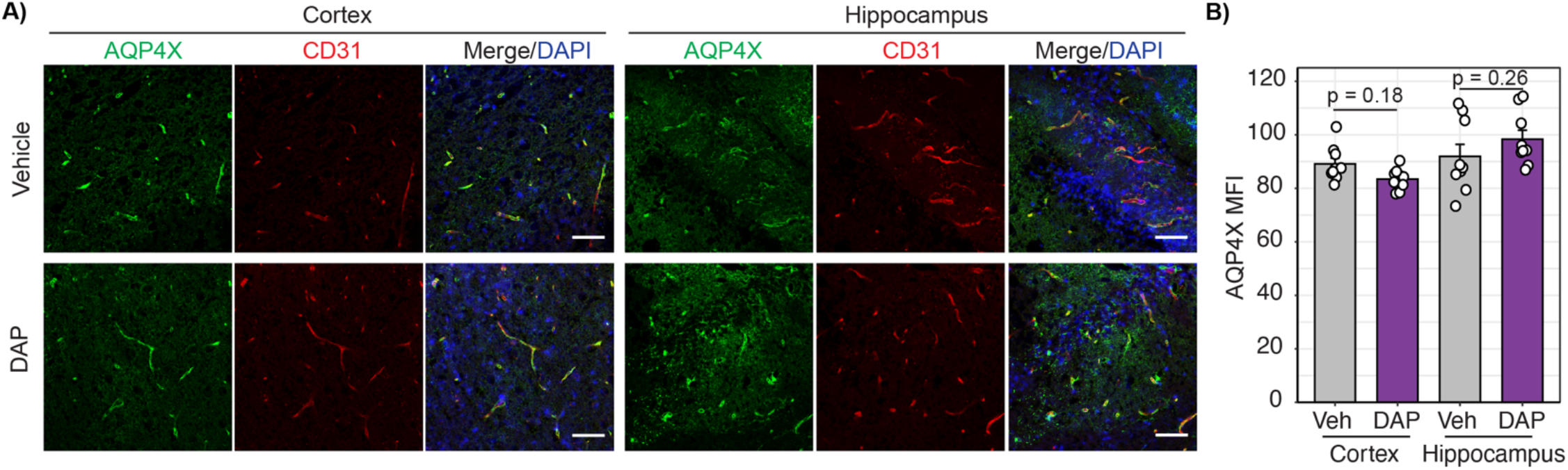
DAP elevates AQP4X levels via translational readthrough. **(A)** Aqp4AllX+/+ mouse brain hemispheres injected with DAP (300 uM) or vehicle, and AQP4X and CD31 visualized in the cortex and hippocampus at or near the injection site using immunofluorescence staining after 20 hours. DAPI marks nuclei. Scale bars = 50 uM **(B)** Quantification of A for mean fluorescence intensity (MFI) of AQP4X. Student’s *t*-test, N = 3 hemispheres per treatment group. Error bars = standard errors of mean. Circles represent technical replicates.

## DISCUSSION

We report that DAP increases AQP4X abundance, consistent with the promotion of *Aqp4* translational readthrough. For more than two decades, AQP4 has been known to be enriched at the blood-brain barrier and to exist as M1 and M23 isoforms^18^. M1 and M23 arise from translation initiation occurring at two AUGs that lie 69 nucleotides apart. Recently, we and others demonstrated that stop codon readthrough results in M1X and M23X (collectively AQP4X)^16,17^. Because AQP4X comprises the perivascular AQP4, this new insight provided a means to target this specific pool genetically. Based on the present study, it is possible to enhance AQP4X pharmacologically as well.

Perivascular AQP4 plays a key role in brain homeostasis and is strongly implicated in neurological diseases. It drives the clearance of protein aggregates, such as those associated with Alzheimer’s and Parkinson’s diseases, by facilitating CSF perfusion of the brain^11,13,14,29^. The same pool, in turn, is diminished in these diseases, suggesting a vicious cycle that could be interrupted by enhancing or restoring AQP4X. In addition, perivascular AQP4 is the primary target of autoantibodies in neuromyelitis optica^15^. Although enhancement of AQP4X may initially appear counterintuitive in neuromyelitis optica, this possibility warrants a careful investigation because Aqp4AllX mice, which constitutively express AQP4X, exhibit reduced levels of AQP4 orthogonal array particles, the actual antigenic structures attacked by autoantibodies^19^. Overall, this manuscript builds a case for preclinical studies on DAP or similar readthrough enhancing compounds as an intervention for multiple conditions through the modulation of perivascular AQP4.

We show that DAP-mediated AQP4X upregulation is readily detectable in transfected cells and wild-type mice, i.e. when the *Aqp4* transcript supports readthrough at the baseline rate of 20%. In mutant mice that already allow readthrough at 100%, DAP is no longer effective, suggesting it works by enhancing *Aqp4* readthrough. This mechanism is further supported by previous studies proposing DAP as a rescuer of nonsense UGA^21^ and the fact that *Aqp4* stop codon is a UGA. The drug has been shown to inhibit tRNA 2′-O-methyltransferase (FTSJ1), an enzyme responsible for methylating the cytosine 34 of tRNA^Trp^ in the anticodon loop^21^. When tRNA^Trp^ is hypomethylated at this wobble position, it recognizes UGA and allows ribosomes to insert a Trp residue. Interestingly, an Arg residue was detected in place of *Aqp4* UGA in our previous study using mass spectrometry on the wild-type mouse brain^16^. Even if DAP forces a Trp in lieu of Arg, it is clear from the current data that this substitution does not impair AQP4X’s perivascular localization.

Because of its specificity for UGA, DAP is less toxic than classical aminoglycosides such as gentamicin, which enhance readthrough globally by binding to and reducing the fidelity of ribosomes and thus allowing several near-cognate tRNAs to compete with release factors at all three stop codons^30^. DAP was nontoxic to HeLa cells at 25 µM or to mice fed with 1 mg for 5 weeks, and treated cells had 775 transcripts altered by ≧2 folds but no aberrant profile in proteomics compared to DMSO, leading authors to argue the drug is safe^21^. In our hands, the EC50 dose of about 30 µM was not toxic to cultured cells, whereas consistent with prior studies^31^, the maximum dose of 300 µM compromised cell growth. Reasonable doses in dogs (8 or 20 mg/kg daily, IV) were also found to be not lethal, although very high doses led to death in dogs (40 or 50 mg/kg, IV, daily) as well as rodents (≥500mg/kg, IP)^32^. In light of high nephro- and oto-toxicity aminoglycosides and debatable performance of other readthrough enhancers such as atlurin/PTC124^33–36^, DAP may emerge as a candidate for clinical trials following further pharmacological characterization and possibly structural modifications.

Nevertheless, DAP’s target FTSJ1 transfers methyl group to not only tRNA^Trp^ but several other tRNA species^37^. Moreover, in our dual luciferase assay for a *Dmd* UGA nonsense mutation, DAP enhanced readthrough (data not shown). While this is encouraging, as the potential of DAP in Duchenne muscular dystrophy remains unexplored, it also suggests the drug acts on UGA stop codons globally. Thus, DAP’s impact on overall translational fidelity, proteome, and cellular stress must be investigated thoroughly. Unfortunately, antibodies and other reagents are not readily available for peptides encoded by the 3’UTR. High-sensitivity omics and rigorous cell and molecular biology approaches seem fitting. Reported as a UGA nonsense corrector in 2020, DAP is not a failed candidate yet. In preclinical studies, it has restored full-length proteins in cellular and animal models for pathogenic mutations of TP53 and CFTR^21–23^.

The use of intracranial injection is a major limitation of current manuscript. Although it yielded proof-of-principle data, it is not a translationally viable route. Nor does it allow glymphatic or neuromyelitis optica autoreactivity assays for probing the functional outcomes of DAP treatment. Yet, to the best of our knowledge, this is the first study to report an *in vivo* upregulation of perivascular AQP4 through a small molecule. We believe pharmacological elevation of AQP4X is unlikely to exert untoward effects as Aqp4AllX mice— which overexpress the isoform in the CNS, kidneys, retina and other AQP4-positive organs— have normal health and lifespan. If pharmacokinetic and toxicology studies ensure safety and potency, DAP, or other readthrough modulators, could be promising for curbing multiple neurological disorders via perivascular AQP4.

## ACKNOWLEDGEMENTS

We thank the University of Texas at Dallas Imaging and Histology Core for confocal imaging and members of our lab for insightful discussions. This work was supported by the NIH grants R00AG061231 and R01AG093941 and The University of Texas at Dallas Rockford Draper Early Career Development Award to DS.

## METHODS

### Animals

Six to eight weeks old C57BL/6J wild-type (JAX # 000664) and *Aqp4* constitutive readthrough mice (Aqp4AllX, maintained on the C57BL/6J background and reported previously^19^) were used for *in vivo* studies. Aqp4AllX genotyping was performed on postnatal day 7 pups and confirmed at the time of experiment using a custom TaqMan SNP Assay (Thermo Fisher Scientific # 4332077). All animal experiments were approved by the University of Texas at Dallas Institutional Animal Care and Use Committee and performed in compliance with the Association for Assessment and Accreditation of Laboratory Animal Care International guidelines to ensure humane care and minimize discomfort. Mice of both sexes were used. Heterozygote crosses were used to generate the homozygote experimental mice. Animals were housed by line and sex at weaning, with *ad libitum* food and 12/12-hour light/dark cycle.

### Cell culture

Mouse neuroblastoma cell line N2a was used for the dual-luciferase and other *in vitro* studies. Briefly, N2a cells were sub-cultured using DMEM-High Glucose media (Hyclone #SH30243) with added 10% Fetal Bovine Serum (Corning #35010CV) and 1% Penicillin-streptomycin (Invitrogen #15070063) at 37°C, 95% air and 5% CO2. 1X PBS and 0.05% Trypsin-EDTA (Gibco #25300120) were used to harvest the confluent cells from culture flasks and cells were counted using an automated-cell counter (RWD #C100).

### Dual luciferase reporter assay and quantification of readthrough rate and EC50

pSGDluc, a dual-luciferase vector containing Renilla and Firefly, was used (Addgene #119760)^38^. An *Aqp4* test cassette was cloned downstream of Renilla and upstream of Firefly, in frame with the two enzymes. The cassette consisted of the last 15 nucleotides of the coding sequence + stop codon + 81 nucleotides following the stop codon: GTATTGTCTTCCGTA<u>TGA</u>CTAGAGGACAGCACTGAAGGCAGAAGAGACTCCCTAGACCTGGCCTC AGATTTCCTGCCACCCATTAAGGAAACAGATTTGTTA. In a positive control, the TGA stop codon was mutated to TGG.

On day one of assay, the test and control plasmids (150 ng/well) were reverse-transfected into N2a cells (2 x 10^4^ cells/well) in a 96-well culture plate using Lipofectamine 3000 (Invitrogen #L3000015) in opti-MEM reduced serum media (Gibco #31985070) and incubated at 37°C. After 6 h, the transfection media was replaced with complete medium containing vehicle (0.2% DMSO), G418 (Invitrogen #10131027) at 500 ug/mL, or DAP (Sigma #247847) at 1 µM, 3 µM, 5 µM, 10 µM, 30 µM, 50 µM, 100 µM and 300 µM. Treatment was allowed for the next 24 hours. On day two, the Renilla and Firefly luciferase activities were quantified using a Single Tube Assay Kit (Biotium #30081-2) and Synergy HTX luminescence reader with dual injectors (Agilent BioTek) following manufacturer’s instructions. Readthrough rate was calculated as:

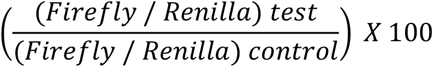

For half-maximal effective concentration (EC50), dose-response data were analyzed in R using the drc package^39^. A four-parameter (slope, lower asymptote, upper asymptote, EC50) log-logistic model was fitted to the data, excluding the 0 µM dose. EC50 was estimated from the fitted curve using the effective dose function.

### Stereotaxical injection

Wild-type and Aqp4AllX homozygote mice were injected intracerebrally using a stereotax fitted with a digital injector (RWD #68537) and a 26-gauge syringe (701, Hamilton). Mice were anesthetized with ketamine/xylazine (100 mg/kg and 10 mg/kg, intraperitoneal) and fixed on the stereotax after shaving the scalp hair. Eyes were lubricated with petroleum jelly, and body temperature was maintained using a heating pad and rectal thermometer. The scalp was cleaned with betadine, ethanol, and saline; locally anesthetized with lidocaine (100 µL of 0.25%, subdermal), and an incision was made along the midline. The bregma and lambda were visualized by swabbing the skull with 3% H202.

For immunofluorescence staining, 0.5 mm burr holes were drilled at anteroposterior (AP) −2 mm, and mediolateral (ML) 1.8 mm relative to the bregma, with a depth of 1.8 mm. The contralateral side was drilled at ML −1.8 mm. 300 µM DAP in a volume of 2 µL was injected into one hemisphere, and equal volume of PBS injected into the contralateral hemisphere. For Western blot, two adjacent sites per hemisphere were administered DAP (400 µM, total volume 3µL) or equal volume of PBS. The coordinates were AP −1 mm and −2 mm and ML 1.8mm. After 10 min, the needles were retracted, and animals were sutured and allowed to recover under observation for 20 hours before harvesting the tissue.

### Western blot

For quantifying the effect of DAP on AQP4X in cell culture, pEGFPc1 plasmid expressing Aqp4 CDS-stop-readthrough region was reverse-transfected into N2a cells (1.5 μg/well, 1X 10^6^ cells/well) in a 6-well culture plate using Lipofectamine 3000 (Invitrogen #L3000015) in opti-MEM (Gibco #31985070). After 37°C at 6 hours, the transfection media was replaced with the complete medium containing vehicle (0.2% DMSO), 500 μg/mL of G418 or 30 µM of DAP. The cells were then continued to be incubated for the next 24 hours before extracting total protein with RIPA buffer (Thermo #87788) with added protease inhibitor cocktail (Roche #11836170001). The proteins were quantified using BCA assay kit (Pierce #2325). For the DAP effect on AQP4X *in vivo*, the mouse brains were harvested 20 hours after stereotaxic injections as described above. The two cerebrums were separated, and the total protein extracted and quantified similarly.

Protein extracts were electrophoresed using 12% Tris Glycine polyacrylamide gels (MP Biomedicals #08W00003) in a mini-Protean apparatus (Bio-Rad #1658004) and transferred onto PVDF membranes using the power blotter-XL system (Invitrogen #PB0013). The membranes blocked in 5% nonfat milk in TBST for 1 hour and incubated overnight at 4°C with anti-AQP4X (1:1000 dilution, rabbit, Cell Signaling #60789S) or anti-AQP4 (1:1000 dilution, rabbit, Cell Signaling 59678S), followed by anti-beta-actin (1:5000 dilution, mouse, Abclonal #AC004). Next, the membranes were washed three times with 1X TBST and incubated with anti-rabbit (Cell Signaling #7074S) or anti-mouse (Cell Signaling #7076S) HRP-linked secondary antibodies at room temperature for 1 hour. Finally, after three washes with TBST again, the membranes were visualized with iBright imager (Invitrogen) using Femto-Plus ECL Substrate (MP Biomedicals #08X100003).

### Immunofluorescence staining

Brains were harvested from stereotaxically injected mice, fixed overnight in 4% paraformaldehyde and dehydrated successively with 10%, 20% and 30% sucrose in PBS at 4°C. After embedding brains in OCT, 14 µM thick sections containing the injection line and 5 sections before and after the injection line were collected on slides using a cryostat (RWD #FS800). Slides were washed twice with 1X PBS, blocked with a buffer containing 5% normal donkey serum and 0.3% Triton X in PBS for one hour at room temperature, and incubated overnight at 4°C with anti-AQP4X (1:200, goat, Shikhar Biotech, custom made against an AQP4 readthrough peptide and knockout-validated) and anti-CD31 (1:50, rat, BD Biosciences #550274) diluted in blocking buffer. Next day, sections were washed with 1XPBS three times and incubated with donkey anti-goat (1:500, SouthernBiotech #642030) and donkey anti-rat (1:500, Invitrogen #A21209) secondary antibodies diluted in blocking buffer for one hour at room temperature. After the same washes, sections were stained with DAPI (Sigma #MBD0015) for 10 min and mounted with ProLong Gold antifade medium (CST # 9071S) for microscopy. 40X images of the cortex and the hippocampus were captured using Leica SP8 confocal, keeping exposure parameters constant across slides.

### Image quantification and statistical tests

ImageJ (NIH) was used for signal quantification in 8-bit images. For densitometry of Western blot, bands were selected using identical-sized rectangles, and integrated densities were obtained. No background subtraction was applied. For mean fluorescence intensity, AQP4X channels were thresheld, keeping parameters constant across images, and area, integrated density, mean gray value limited to thresheld areas were obtained using a macro. Both total and perivascular AQP4X were quantified, the latter using CD31 as a mask. Owing to the perivascular localization of AQP4X, the two values yielded identical results, and only total mean fluorescence intensities were shown in the manuscript. Researchers involved in quantification were blind to treatment paradigms.

One-way ANOVA followed by Dunnette’s post-hoc was used to compare three groups (Fig. 1). Unpaired two-tailed Student’s *t*-test was used to compare two groups (Figs. 2 and 3) although paired t test was also applicable given the drug and vehicle treatments occurred in the same brain. Assumptions for normality, homogeneity of variance, continuous scale tests were ensured in R while applying the tests. Only those animals with suboptimal stereotaxic surgery were excluded from analysis. Plots wre generated with ggplot package. Bar plots show mean ± standard error of the mean and individual data points. Box plots show minimum, maximum, quartiles, median and outliers. Technical replicates were averaged to yield a single biological replicate. Sample sizes were informed by prior reports^21,22^ and pilot experiments, but not by a priori power calculation. A *p* value < 0.05 was considered statistically significant.

